# Climate change and deforestation boost post-fire grass invasion of Amazonian forests

**DOI:** 10.1101/827196

**Authors:** Bruno L. De Faria, Arie Staal, Philip A. Martin, Prajjwal K. Panday, Andrea D. Castanho, Vinícius L. Dantas

## Abstract

Interactions among climate change, deforestation and fires are changing the stability of the Amazon forest, and may promote transitions to degraded grassy ecosystem states. However, our ability to predict the locations in the Amazon that are most vulnerable to these transitions is limited. In this study we used a dynamic carbon model to evaluate how drought, climate change and deforestation could affect the probability of post-fire grass invasion across the Amazon, and identify where grass-fire feedbacks may promote the persistence of species-poor degraded forests with savanna-like structure. Our results suggest that, under current climatic conditions, post-fire grass invasion could affect 11% of the Amazon, with the south-eastern Amazon at highest risk of invasion. We forecast that under business as usual climate change, by the end of the century areas with a high probability of post-fire grass invasion will increase to 20% of the Amazon. In 10% of the Amazon fire return interval will be shorter than the time required for canopy recovery, implying high risk of irreversible shifts to a fire-maintained degraded ecosystem state. Although resilience in canopy regeneration is evident in areas with low fire frequency, increased fire frequency could inhibit regeneration even in forests where grass is currently excluded, and push the Amazon forests towards a tipping point causing large areas of forest to transition to low tree cover state.

## INTRODUCTION

Tropical forests contain between half and two thirds of terrestrial global biodiversity and provide vital ecosystem services at local, regional, and global scales (Dixon et al. 1994; Foley et al. 2007; Marengo et al. 2018). However, these forests are undergoing widespread loss and fragmentation as a result of deforestation as well as degradation from climate change and fire (Hansen et al. 2013; Espírito-Santo et al., 2014). Forests in the tropics are expected to be especially sensitive to such changes. Modeling (Hirota, et al. 2011; Van Nes et al. 2018), observational (Dantas, Batalha, & Pausas, 2013; Dantas et al. 2016) and experimental (Silverio et al. 2013) studies suggest that a positive feedback between forest loss and fire may cause a shift of closed canopy forest to a grassy savanna-like ecosystem. These novel ecosystems are predicted to differ dramatically in species composition, contain fewer species, and especially fewer endemic species, than both old growth savanna and forest (Veldman & Putz 2011; Veldman et al. 2015). However, at present, it is unclear how the local process of post-fire grass invasion, increasing forest vulnerability to grass-fire feedbacks, could scale-up to affect large forest regions.

One forest region that may face the threat of such a shift to a degraded savanna-like state is the Amazon. This threat is partly driven by a recent sharp increase in fire frequency (from once every 500–1000 years, prior to modern-day human colonization, to once 5-10 years; Barlow & Peres 2008), due to deforestation and climate change (Balch et al. 2015; Fearnside 2013; Gutiérrez-Vélez et al. 2014). In addition to fire frequency, climate change is predicted to increase fire intensity as a result of reductions in annual rainfall and increases in temperature. These changes in regional climate are expected to increase water deficits reducing litter moisture contents and increasing litter quantity, which will likely promote fires of much higher intensities (De Faria et al. 2017). Because single high-intensity fires can cause high damage to above-ground plant parts in tropical forests (Brando et al. 2012, 2014), climate change could drive higher forest canopy cover losses after fire events by increasing fire intensity.

In addition to climate change, forest loss and fragmentation may magnify losses caused by fires, as evidence suggests that canopy cover can be reduced by up to 60% in areas within 3 km of a forest edge (Broadbent et al. 2008; Wuyts et al. 2017). Combined, the effects of climate change on fire intensity and deforestation in edge areas are expected to result in very high canopy cover losses in the region. As a result, the risk of reaching a critical canopy cover threshold beyond which highly flammable shade-intolerant grasses can establish in the forest understory is expected to increase. (Hoffmann et al. 2012; Silvério et al. 2013; Cardoso et al. 2018). Once this threshold is crossed, both fire intensity and frequency are likely to increase abruptly, as the high cover by low bulk density grass fuels in the understory dramatically increases fuel flammability (Hoffmann et al. 2011). This could potentially create positive fire-grass feedbacks that could cause regime shifts to low tree cover states (Hoffmann et al. 2009; Staver, et al. 2011; Dantas et al. 2016; Van Nes et al. 2018; Staal et al. 2018a). Grass invasion appears to be higher in highly fragmented forests (Balch et al. 2015) and increases in fire intensity resulting from climate change are expected to greatly expand these areas. To avoid being arrested in a fire trap (Murphy et al. 2010; Grady & Hoffman 2012; Trauernicht et al. 2016), the forest must be resilient, that is, recover canopy cover quickly enough to exclude shade-intolerant flammable grasses before the next fire (Hoffmann et al. 2012). Many studies in both Africa and South America have identified a Leaf Area Index (LAI) of 3 as the critical canopy cover threshold at which shade-intolerant C4 grasses are excluded from the forest understory (Hoffmann et al. 2012, Dantas et al. 2016, Cardoso et al. 2018). How climate change and deforestation will affect post-fire grass invasion and the resilience of forests to grass-fire feedback is not currently known.

We simulated grass invasion as a function of canopy cover losses after a fire event (as a function of how climate affects fire intensity) and from logging (within forest edge areas) for a fire taking place under present and future climatic conditions in the Amazon region. We also evaluated the reversibility of grass invasion by contrasting simulated vegetation recovery time (as a function of climate) and fire return interval of a given location. Using this framework, we address the following questions:

1. Where in the Amazon are forests most vulnerable to post-fire grass invasion?
2. How will predicted increases in fire intensity under climate change and deforestation affect these areas?
3. Where in the Amazon are the least resilient forests that are most likely to shift to a novel ecosystem state with savanna-like structure, under present and future conditions?

We hypothesized that the already drier climate and higher deforestation rates in the south-eastern Amazon (Silva-Junior et al. 2018) would result in high probability of grass invasion and increased fire frequencies in this part of the basin. Since the south-eastern region is also predicted to experience increases in temperature and decreases in precipitation under climate change (Chen et al. 2011; De Faria et al. 2017; Phillips et al. 2009), we hypothesized that grass invasion will largely increase in the region, undermining forest resilience to state shifts.

## METHODOLOGY

Our study focuses on the Amazonia which contains approximately 5.5 million km^2^ of tropical forest. For this study, we used a process-based carbon cycling model, CARLUC-Fire (De Faria et al. 2017) to simulate the probability of post-fire grass invasion following fire in the Amazon. We used this model to investigate vulnerability to post-fire grass invasion, both under the current climate and under business-as-usual climate change. Below we outline our study design in detail.

### The model

The CARLUC-Fire model is a modified version of the Carbon and Land Use Change dynamic carbon model CARLUC (Hirsch et al. 2004) which incorporates forest flammability, fire behaviour, and the impacts of fire (De Faria et al. 2017). The CARLUC is a process-based, spatially-explicit model and estimates net primary productivity (NPP) based on microclimatic conditions as well as litter, woody debris, and humus resulting from plant mortality (for more detail see supplementary materials). Each time step of the model represents a month.

The CARLUC-Fire model (De Faria et al. 2017) specifically accounts for the effects of fire by estimating forest carbon losses after a fire as a function of fire intensity. CARLUC-Fire uses two loss terms: drought-induced loss of above-ground live carbon stocks and fire-induced loss mediated by changes in fuel loads and moisture. These losses are connected because the former contributes to fuel loads, affecting fire intensity. Thus, climate modulates forest carbon both directly (drought-driven branch and leaf loss) and indirectly (fire-induced post-fire plant mortality, resulting from climate-mediated changes in fire intensity). In this study we adapted the model to predict changes in canopy cover as a function of fire intensity and the associated probability of grass invasion, and to simulate canopy recovery after a fire (see supplementary materials for details).

### Experimental runs

We ran our simulation under two scenarios: one in which climate mirrored current conditions and one in which climate mirrored a business-as-usual predictions for climate change for the period 2070-2099.

To determine the probability of grass invasion after a fire under current conditions, we ran the model using mean climate conditions for 1980-2009. Climatic conditions were based on the Climatic Research Unit dataset (CRU TS; Harris et al., 2014) and the Integrated Biosphere Simulator (IBIS) dynamic vegetation model, which was forced with the same CRU climate data (Panday et al 2015). Simulated droughts were parameterized using data from NASA’s Tropical Rainfall Measurement Mission (TRMM, data product 3B43). Pre-fire canopy cover was based on the MODIS LAI product (MCD15A2H; Mynen et al. 2015). We excluded deforested areas using deforestation maps from the annual Landsat-based Project for Monitoring Amazonian Deforestation (PRODES; INPE, 2017) and MapBiomas collection 3. Following this, we used simulations to estimate the probability that a single fire event in a year of drought would promote grass invasion across the Amazon under current climate (see supplementary materials for more details of methods).

To evaluate the potential impacts of climate change on the probability of grass invasion, we ran the simulation using a scenario for 2070-2099 based on a Representative Concentration Pathways (RCP8.5 scenario) of the Coupled Model Intercomparison Project Phase 5 (CMIP5) multi-model ensemble. This scenario assumes a continued increase in greenhouse gas emissions (Duffy et al. 2015), leading to air temperature increasing by approximately 4-5°C across the southern Amazon, and reduced precipitation during the dry season (De Faria et al. 2017; Justino et al. 2011; Phillips et al. 2009). To parameterize this scenario we used air temperature and precipitation projections from 35 climate models participating in the CMIP5.

In addition to the impacts of climate change, we also analyzed the impact of edge effects resulting from logging and forest fragmentation on the probability of grass invasion under current and future conditions. Evidence suggests that edge effects can reach 2-3 km from the border, especially logging, which alone can reduce canopy cover by up to 60 % (Broadbent et al. 2008, Wuyts et al. 2017). To simulate the effects of logging on forest edges we imposed an additional 60% loss in post-fire LAI in areas ≤3 km from the forest border, before calculating the probability of grass invasion.

To examine the effects of uncertainty regarding the impacts of edge effects on LAI we also performed simulations in which losses of LAI were 10 % rather than 60% loss. The difference between this and the simulation assuming 60 % loss at the edges was used as a measure of the uncertainty associated with this assumption.

Distance to forest edges was calculated as the distance to a deforestation cell using the “gdistance” packagein R (van Etten, 2017). Deforestation and forest cover were used to define forest borders and were defined using data from PRODES with cumulative deforestation up to 2017. Edge distances were not updated after applying fire-induced losses.

To identify locations that may be vulnerable to being trapped in a grass-fire feedback loop we compared the time required for each pixel to recover an LAI value of 3 (thereby allowing for exclusion of grasses) with the observed satellite-derived mean fire interval between 2003-2016. LAI recovery time was calculated as a function of our climate input variables using equations for forest productivity (Hirsch et al. 2004). We considered a forest area to be resilient when the time required to achieve an LAI of 3 was shorter than or equal to the mean fire interval, and non-resilient otherwise. Fire return interval per pixel (FRI) was calculated using MODIS Burned Area Product Collection 6 (MDC64A1; Giglio et al. 2018) and The Global Fire Atlas dataset (Andela et al. 2019). Based on empirical observations for forests and savannas (Dantas et al. 2016) we reduced the mean fire interval by 50% in areas where LAI values drop below 3 following a fire. We did not model changes in FRI resulting from climate change because the relationship between fire probability and climate in South America is non-linear and complex (e.g. Lehmann et al. 2011).

All analyses of simulations were carried out in R (R Development Core Team, 2012) using the packages rgdal (Keitt et al., 2014), raster (Hijmans et al. 2014), and rasterVis (Perpinan & Hijmans 2018).

## RESULTS

We find that under the current climate (1980-2009), the areas which would show the highest probability of grass invasion if affected by fire are located in the southeastern part and, on a smaller scale, in the southwestern part of the Brazilian Amazon (Acre state) (Fig. 1). These areas cover 560,000 km^2^, approximately 11% of the total forest area (Fig. 1A). However, under business-as-usual climate change, the total area with high probability of grass invasion increases to 1.0 million km^2^ by the end of the century (2070-2099), a 78% rise. This area amounts to 20% of Amazonia, most of which is in the southeastern portion of the basin (Fig. 1B). These levels of grass invasion would result in strong changes in the frequency distribution of LAI, especially under climate change. This results from shifts in local forest LAI (as well as regional-scale modal LAI) towards lower values, increasing the area with an LAI below 3 (Figs. 2 and S3). Climate change alone has a very subtle effect on forest productivity as simulated by the IBIS model (Fig. S3), and, thus, the changes under climate change were mainly explained by changes in fire intensity. Fires increase the area with high probability of grass invasion (LAI < 3) by only 6 % under current conditions, much lower than the 80 % increase under future conditions.

**Figure 1.**
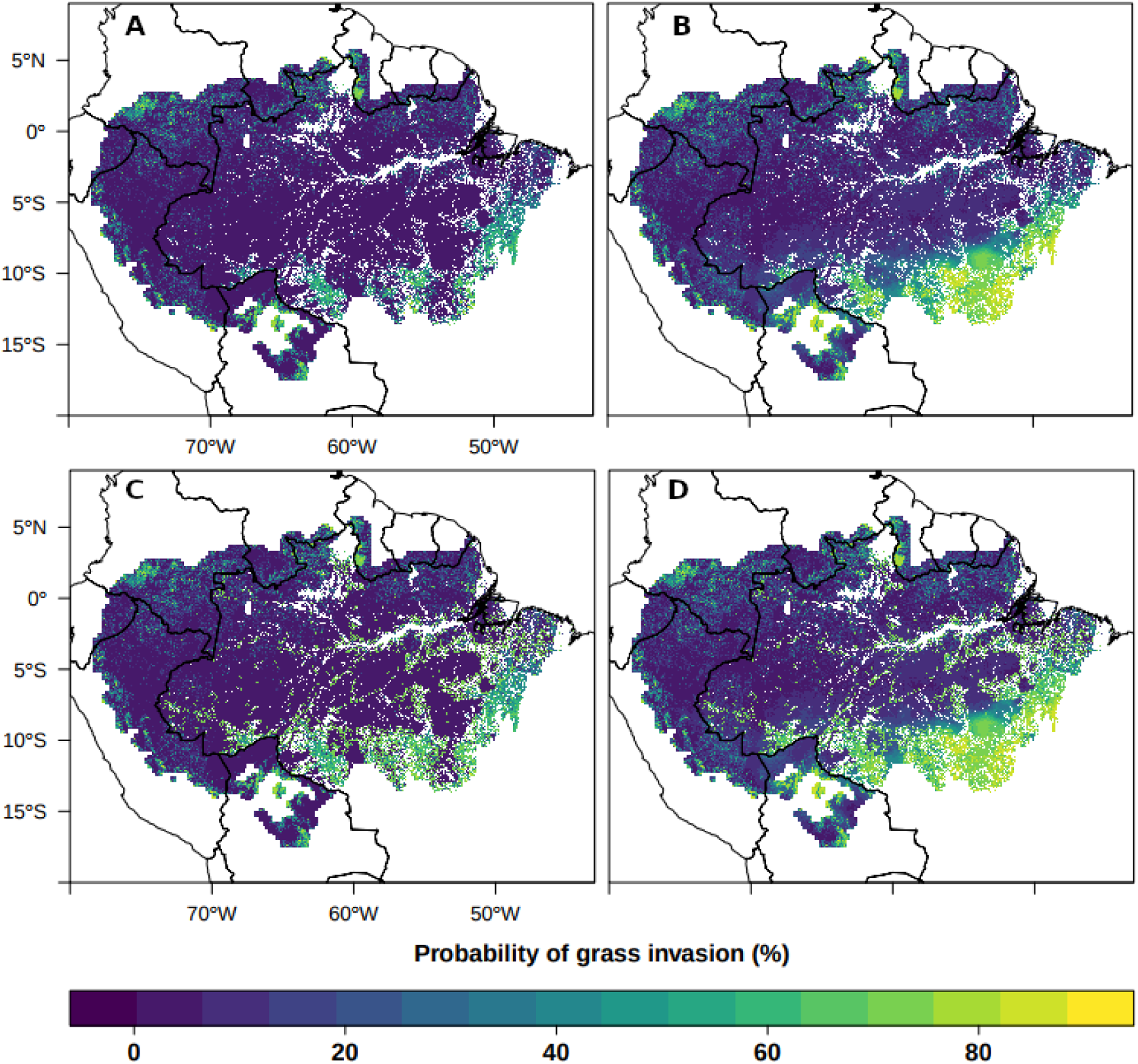
Probability of grass invasion (%) after fire calculated based on post-fire Leaf Area Index (LAI) and edge effects resulting from deforestation. (A) The probability of grass invasion under current (1980-2009) conditions. (B) The probability of grass invasion under average conditions projected for 2070-2099, in a business-as-usual climate change scenario. (C-D) The probability of grass invasion after fire in A and B, respectively, including edge effects. Our simulations assume fire to take place in a drought year.

**Figure 2.**
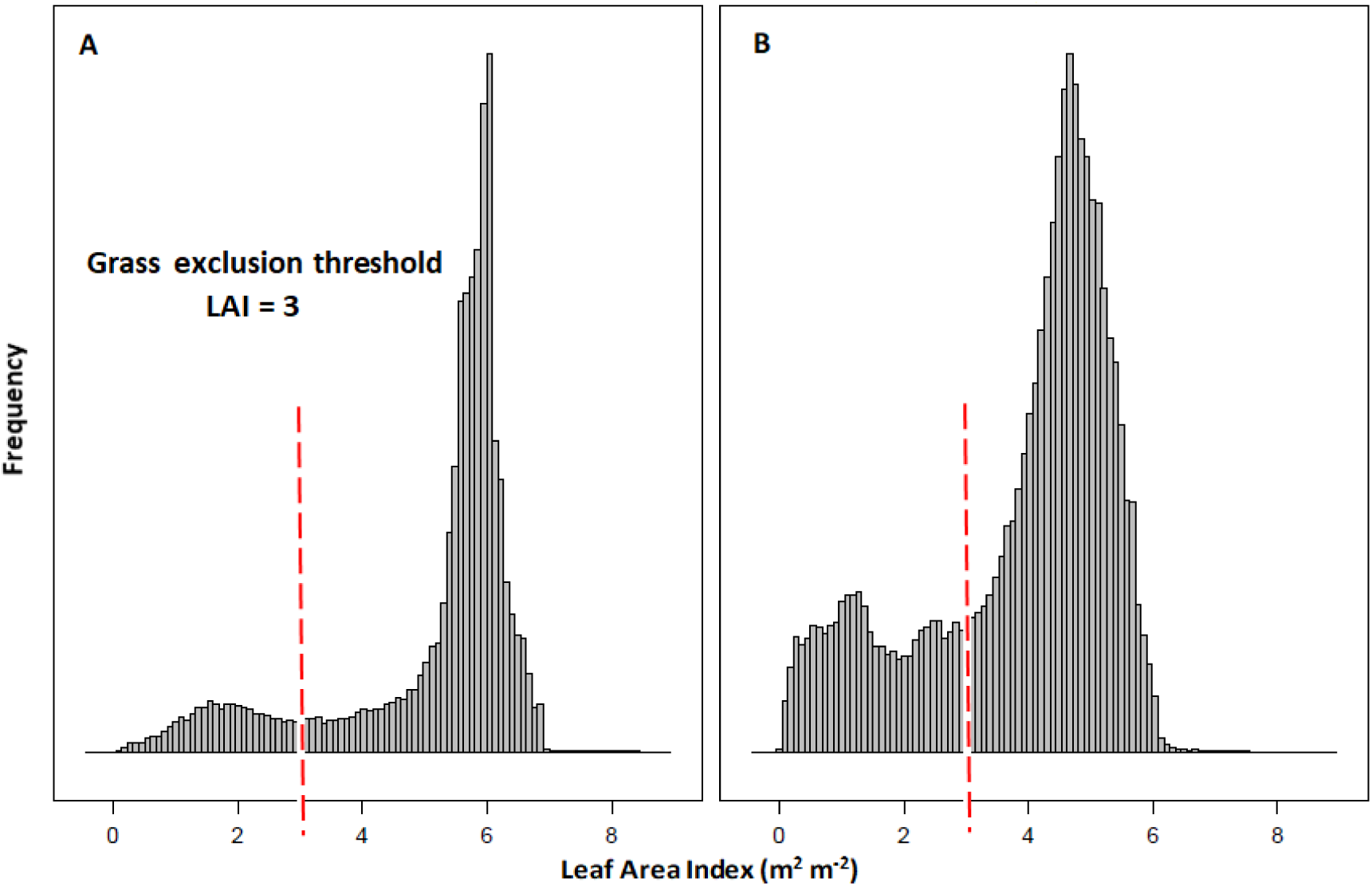
Frequency distributions of Leaf Area Index (LAI) after a fire for the Amazon region under current (A) and future (B) climate scenarios. Red dashed lines indicate the grass exclusion threshold, the critical LAI value (LAI = 3) above which the forest has sufficient canopy cover to exclude shade-intolerant grasses.

Approximately 180,000 km^2^ of forest patches are within 3 km from a forest edge (Fig. 1C). Accounting for edge effects resulted in an increase in the areas under high risk of grass invasion by 30 %, totalling 740,000 km^2^ (Fig. 1C) and 1.13 million km^2^ under current and future conditions, respectively (Fig. 1D). In case of the more conservative scenario in which canopy cover losses in edge areas are only 10 % instead of the assumed 60 %, grass invasion probability is 17 % lower.

There are substantial spatial differences in recovery time (post-fire time required to achieve LAI = 3). The southern and southeastern parts of the basin currently require the longest recovery times (mean = 4.6 years; median = 5.1 years). In total, an area of 162,000 km^2^ would require fire intervals longer than five years to achieve an LAI of 3. Climate change doubles this area (Fig. 3B), increasing the area requiring a fire interval longer than 1 year by 52 % in southeastern Amazon (from 659,000 to 1.0 million km^2^). Other areas (244,000 km^2^) are not substantially affected as the impacts of fire are small and, even when LAI fell below 3, recovery could take place within a year, even in the business-as-usual scenario (Figure 3 A, B). Mean fire return intervals in the Amazon range between 1 and 15 years, and is lowest in human dominated areas (FRI of 1-3 years; Fig. S2), where deforestation and human land uses, especially agriculture, are substantial. Increases in the frequency of future fires as a result of grass invasion result in a substantial increase in areas with low fire return interval (Figure S2).

**Figure 3.**
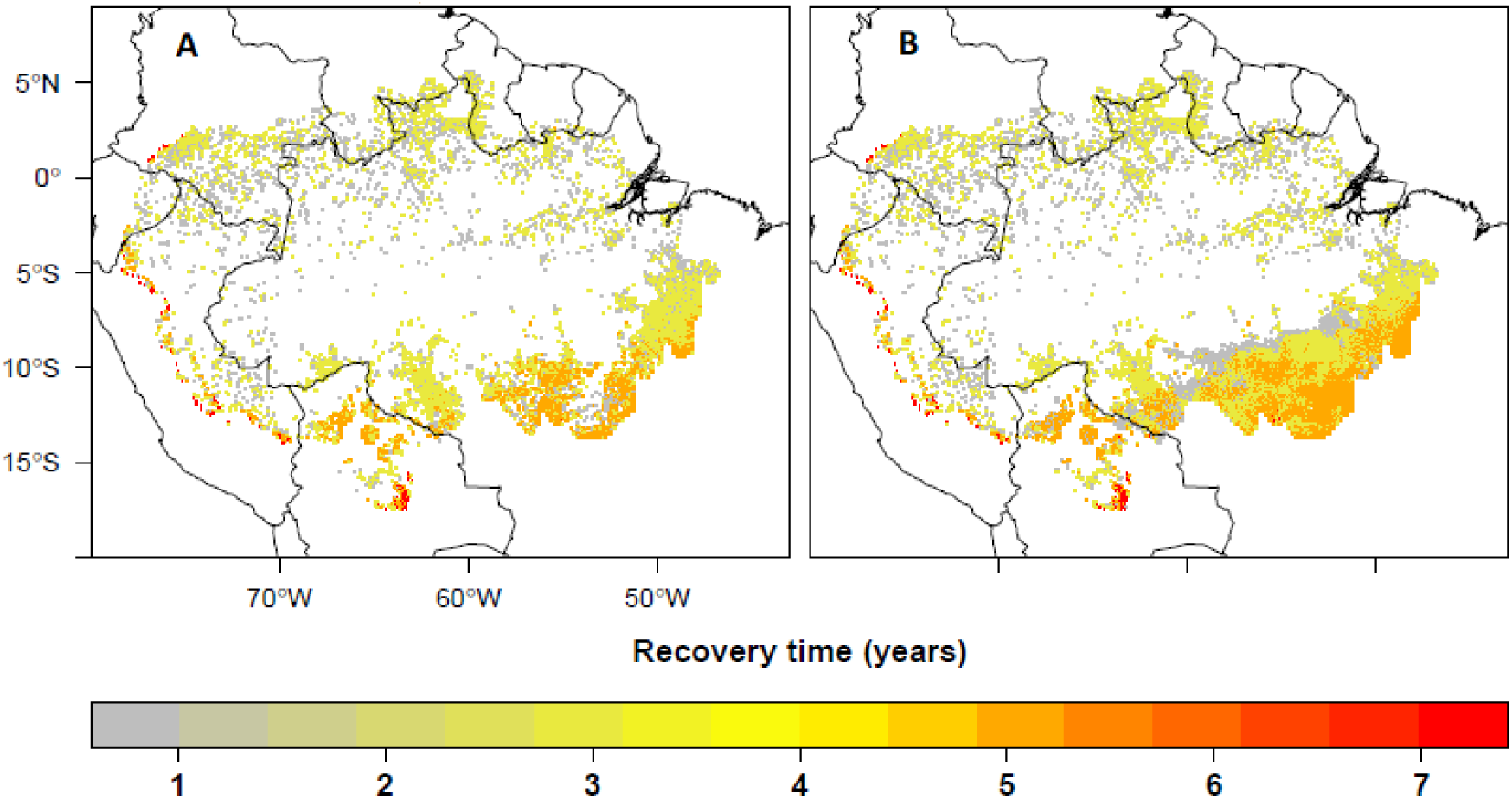
Present (A) and future (B) recovery time required for grass exclusion across the Amazon region. Grass exclusion occurs when the forest develops a critical amount of canopy cover (Leaf Area Index = 3) at which shade-intolerant grasses are outcompeted by forest trees.

Non-resilient areas, where recovery time exceeds fire return interval, could emerge in approximately 167,000 km^2^ under current climate (Fig. 4A), and in about three times this area in the future (449,000 km^2^) (Fig. 4B). This implies that approximately 10 % of the forest in the Amazon basin may be at risk of a regime shift to a low tree cover state by the end of the 21st century.

**Figure 4.**
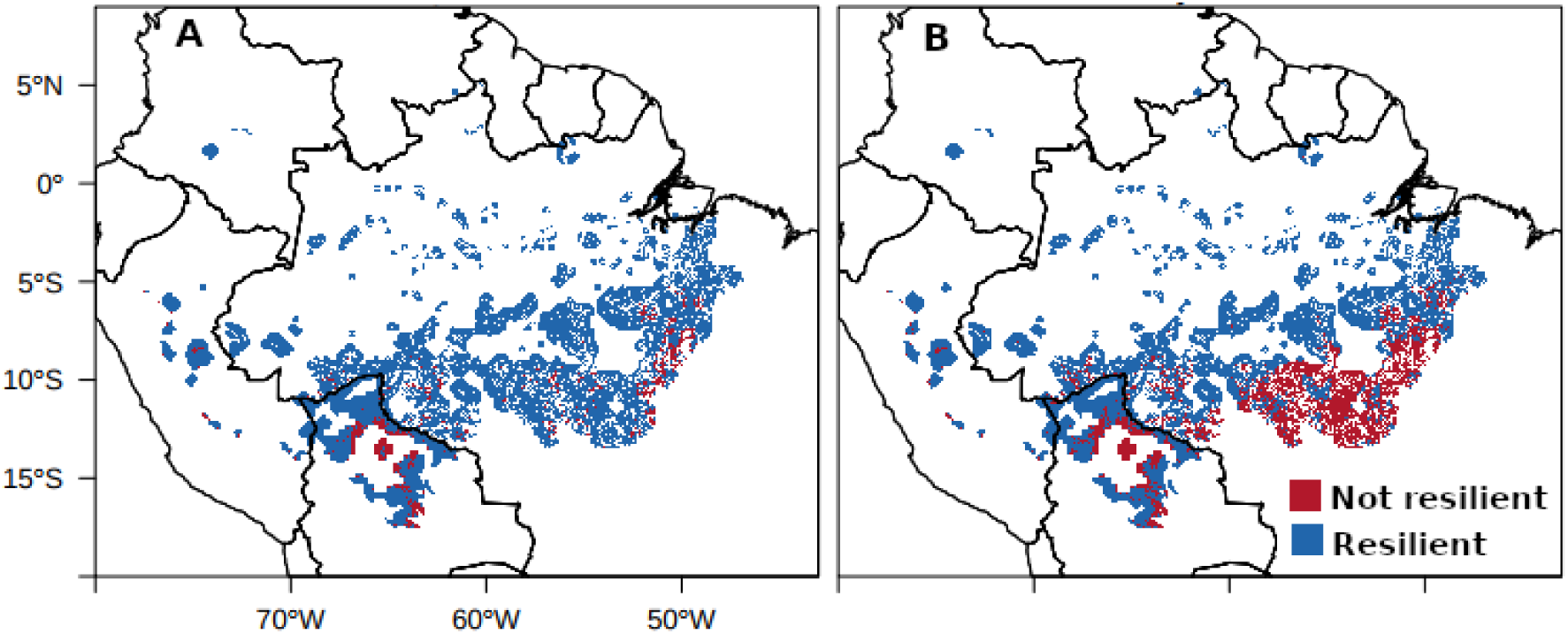
Resilient and non-resilient forest areas under current (A) and business-as-usual climate change (B) conditions. Resilience is based on the difference between the Fire Return Interval (FRI) and forest recovery time. A site is considered resilient (blue) when the time required for the forest to recover an LAI of 3 and exclude shade-intolerant grasses is shorter than the FRI, and not resilient (red) otherwise.

## DISCUSSION

Our model predicted that post-fire grass invasion could affect 11 % of the Amazon under current climatic conditions, which under business-as-usual climate change could rise to 20 % by the end of the century. This implies a dramatic increase in both fire recurrence and intensity in the case a subsequent fire precedes total canopy recovery. The predicted increase under climate change is mostly due to increased fire intensities under future drier and hotter climates, driving greater post-fire canopy cover losses. In both current and future climate, the areas with a high probability of post-fire grass invasion were predicted to be concentrated in the southeastern Amazon, which is consistent with previous empirical studies showing that grass invasion after fire already affects some of these areas (Veldman et al. 2009, Balch et al. 2015). We also found that 3 % of the forest patches in the Amazon were located within 3 km from an edge, most of which was concentrated in the southeastern region. Edge effects resulting from deforestation were predicted to increase the area affected by post-fire grass invasion from 560,000 to 740,000 km^2^ under present conditions and from 1.0 million to 1.13 million km^2^ under future conditions, totalling 14 and 23 % of the region. Therefore, we expect that both climate change and deforestation will greatly increase grass invasion in the southeastern Amazon.

Many locations with a high probability of grass invasion currently experience recurrent fires. This includes areas that are predicted to show the longest recovery periods, many of which require an interval longer than 5 years. This results in a high probability that a subsequent very-high-intensity grass-fueled fire would occur before grass exclusion, driving even larger decreases in LAI (Hoffmann et al. 2012, Silvério et al. 2013, Dantas et al. 2016). This could lock the ecosystem in a feedback loop, in which fire promotes grasses by reducing tree cover and grasses produce frequent and intense fires (Warman and Moles 2009, Brando et al. 2010, Pausas and Dantas 2017). As a result, the ecosystem would be maintained in a savanna-like state, but with much lower carbon stocks and many fewer species (Veldman and Putz 2011, Veldman et al. 2015) than found in old-growth South American savannas. According to our results, as much as 10 % of the Amazon, approximately 449,000 km^2^ (or 449 million hectares), could experience such shifts in the future, which would likely be very difficult to reverse.

The probability of areas being trapped in a grass-fire feedback loop depends on sufficiently frequent fires. Here we used current fire return interval, corrected by increases resulting from grass invasion (see Methods), to predict the areas that could potentially experience irreversible state shifts. However, fire frequencies could also increase in the future due to predicted increases in drought frequency (Aragão et al. 2018). In fact, a recent study predicted that burned area could increase by 39-95 % in the nearby, but more seasonal, savanna region of Brazil, as a result of climate change (Silva et al. 2019). The extent to which the Amazon will face similar increases is unknown. In addition, several other geophysical feedbacks can take place after grass invasion that could increase fire frequency and preclude a fast forest recovery (Archibald et al. 2018). For instance, canopy cover losses are expected to positively feed back to drought (Staal et al. 2018b), further increasing ecosystem flammability (Staver et al. 2011). Moreover, increased fire frequency is associated with long-term soil nutrient losses (Pellegrini et al. 2018), which could decrease the ability of the vegetation to recover before the next fire (Flores et al. 2019). Synergisms with other disturbances than fire, such as windstorm, could greatly amplify fire-driven mortality and affect recovery, especially in edge areas (Silverio et al. 2017; Brando et al. 2019). Finally, tree regeneration is expected to become much slower once grasses have invaded due to the competition between tree seedlings and grasses (February et al. 2013; Hooper, Legendre, & Condit, 2005), and frequent fires in grass-dominated ecosystems have been shown to reduce tree growth up to 66 % (Murphy et al. 2010). Predicted increases in atmospheric CO_2_ concentrations are unlikely to counteract these effects, in part due to the already nutrient-limited nature of most tropical soils (Ellsworth et al. 2017). Thus, the area with potential for transitions to alternative ecosystem states could be larger than the conservative estimates shown here.

Land-use changes may accelerate the rate of canopy cover reduction by: (i) reducing tree cover and creating vulnerable forest edges (Balch, Nepstad, & Curran, 2009, Morton et al. 2013); (ii) introducing grass propagules (Veldman & Putz 2010, 2011); and (iii) increasing ignitions associated with land management practices (Nepstad et al. 1999, Cochrane et al. 1999; Armenteras et al. 2013). Logging activities alone currently taking place in edge areas can reduce canopy cover up to 60 % (Broadbent et al. 2008). As we show here, these activities can dramatically increase grass invasion in these areas. With 3 % of the forests located within 3 km from an edge, the Amazon forest already faces a risk of extensive grass invasion under current climatic conditions, and this amount will greatly increase with climate change. Most of these areas are located in the southeastern Amazon. Predicted increases in land-use changes (Aguiar et al. 2016) could make grass invasion similarly likely across other parts of the Amazon region in the future and even increase the probability of fire feedback loops starting in other locations. In fact, since fire frequency and intensity is usually higher in edge areas (Broadbent et al. 2008, Balch et al. 2015, Silva-Junior et al. 2018), the extension of these areas could be greater than estimated here if future fragmentation and land-use changes were considered. Regarding land-use changes, however, fire frequency does not necessarily increase with rising deforestation rates, as the decrease in fuel connectivity also limits fire spread, depending on land use types, resulting in shorter and less frequent fires at the global scale (Andela et al. 2017; Archibald et al. 2013).

Several of the model parameters were derived from a single study in the southeastern Amazon. Therefore, our model did not take into account local factors such as soil and forest types, management history, and flooding and windstorm regimes, when simulating forest recovery time and fire-induced losses. However, these factors likely affect fire-mediated losses and forest resilience. For instance, seasonally flooded forests are less resilient to fire than upland forests due to the rapid loss of soil fertility (Flores et al. 2017). Moreover, trees growing on nutrient rich clay soil can recover faster than plants on white sand nutrient poor soils, and insect herbivores density modulate this effect (Fine, Mesones, & Coley, 2004). In addition to habitat, forest species traits also vary in space and time, and there are a suite of plant traits that reduce fire (bark thickness, tree size, height, and wood density) and herbivory (e.g. tough and chemically protected leaves) damage, and an additional suite that confer an advantage in postfire recovery (e.g. sprouting capacity, fast growth rate; Fine et al. 2004, Hoffmann et al. 2009, Balch et al. 2015). Explicitly considering these factors could greatly improve the accuracy of our model. Moreover, climate change is likely to direct affect fire frequency, while our model only considered the indirect effects of climate change, that is, mediated by changes in grass cover, on fire frequency. Evidence shows that grass presence is a key determinant of fire frequency (Dantas et al. 2013). However, it is likely that incorporating the direct effects of climate change on fire frequency in our model could increase the precision of the findings. Finally, our simulations assume fire to take place in a drought year, and evidence shows that fires in wetter year may not drive substantial grass invasions. However, drought events are relatively frequent and are predicted to become more frequent and intense under climate change (Duffy et al 2015, Malhi et al 2009).

## CONCLUSION

In this study, we show that large parts of the southern and southeastern Amazon are at risk of post-fire grass invasion. Some of these areas may already experience sufficiently frequent fires to cause a shift to a savanna-like state, and these areas could dramatically increase in response to climate change and fragmentation. Although resilience in canopy regeneration is evident in areas with low fire frequency, increased fire frequency due to climate change, deforestation and fragmentation, as well as the associated feedbacks, could, in many areas, preclude the regeneration of forest cover and push them towards a tipping point, after which they are maintained in a degraded low tree cover state. If such a transition occurred on large scales it would have major impacts for Amazonian biodiversity (Barlow and Peres, 2008), as well as on the ecosystem services provided by the forest at both local and global scales. To avoid these negative impacts, two complementary strategies are required. First, global action to limit greenhouse gas emissions is required in order to reduce the potential for severe climate change. Second, in order to limit anthropogenic fires, we recommend that new protected areas are created, that effective monitoring systems are implemented, and that fire-free agricultural practices are encouraged, especially in the most vulnerable, southeastern, part of the basin.

## ACKNOWLEDGEMENTS

We thank the Federal Institute of Technology North of Minas Gerais (IFNMG-*campus* Pirapora and Diamantina), Federal University of Uberlandia (UFU) and Federal University of Vale do Jequitinhonha e Mucuri (UFVJM). We thank A. R. Rech, M. T. Coe, D. Silvério, P. Brando, L. Rattis, E. Pinagé for support and assistance with this research. AS acknowledges support from the European Research Council project Earth Resilience in the Anthropocene (743080 ERA).

## Author contributions

Conceptualization: BLF, AS, PAM, VLD.

Participated actively in execution of the study: BLF, AS, PAM, VLD.

Analyzed and interpreted the data: BLF, AS, PAM, VLD.

Data curation: BLF, PkP, ADC.

Visualization: BLF, AS, PAM, VLD.

Writing – original draft: BLF, AS, PAM, VLD.

Supervision: VLD.

Writing – review & editing BLF, AS, PAM, PkP, ADC, VLD.

