## SUPPORTING INFORMATION for "Climate change and deforestation boost post-fire grass invasion of Amazonian forests"

### EXTENDED METHODS

#### Model description

CARLUC model estimates net primary productivity (NPP) from plant available water (PAW), photosynthetically active radiation (PAR), vapor pressure deficit (VPD), and air temperature. During each monthly time step, NPP is allocated to wood, leaf, and root carbon pools. Death of plants results in dead organic matter which is allocated to structural leaf litter, metabolic leaf litter, structural root litter, metabolic root litter, coarse woody debris, and humus pools (for details, see Hirsch et al. 2004). We consider leaf litter and small woody fuels (i.e. 1 h fuels) as the fuel load.

CARLUC-Fire (De Faria et al. 2017; Figure S4) is an extension CARLUC (Hirsch et al. 2004) with a fire component. Following De Faria et al. (2017), fire-induced biomass loss is a function of fire intensity. Fire intensity (FI, $kW \cdot m^{-1}$) measures the rate of energy released along the fire front, and is strongly correlated with the above-ground impacts of fire. Given that fire intensity and damage to above-ground plant parts are highly correlated in tropical forests (Brando et al. 2012, 2014), especially ‘topkill’ in woody plants (Higgins, Bond & Trollope 2000), a high fire intensity implies higher canopy cover losses. The intensity of a fire depends on on fire spread rate (FSR, $m \cdot min^{-1}$) and the mass of fuel consumed by fire (W, $kg \cdot m^{-1}$). Both FSR and W depend on litter moisture content (LMC, %), while the latter is also a function of load mass

(Table S1). Fuel conditions and loads are influenced by climate. Fuel moisture declines with increasing temperature and vapor pressure deficit (VPD; Ray et al. 2005), while fuel amounts increase with water stress represented by maximum climatological water deficit (MCWD), mediated by the effects of the difference between precipitation and mean regionwide evapotranspiration on leaf and branch shedding (i.e the relationship between MCWD and changes in biomass (Phillips et al. 2009)). This relationship was derived from the Amazon forest inventory network (RAINFOR). When MCWD drops below −40 mm, this relationship predicts that as water stress increases (represented by MCWD) so do associated losses in aboveground biomass (Eq. S1).

The relationship between fire intensity and fire-induced biomass losses was derived from a large-scale fire experiment in southeast Amazonia (Brando et al. 2014) (Eq. 1). Based on this experiment, percentual loss of Aboveground Biomass (AGB) was calculated as follows:

$$\text{Percent loss of ABG carbon} = 1/(1 + \exp(2.45 - 0.002373 * \text{FI})) \qquad \text{(Eq. 1)}$$

The live carbon pool can be converted into leaf and stem biomass components, and leaf biomass can be used to calculate LAI by multiplying it by a specific leaf area (SLA; the fresh area of a leaf divided by its total mass) value of 20 $m^2$ $kg^{-1}$ (Hirsch et al. 2004). Thus, to calculate LAI loss by fire, we multiplied the carbon loss term of CARLUC-Fire by the proportion of carbon content in leaves (Hirsch et al. 2004) and, then, by the above-mentioned SLA value. We used an empirically derived equation relating LAI and the probability of grass invasion (Silverio et al. 2013) to evaluate grass invasion in each pixel of 3 × 3 km. This logistic function (Eq. S2; Fig. S1) was based on a large-scale fire experiment in southeastern Amazon and captures how changes in canopy cover associated with drought- and fire-induced tree mortality influence the probability of grass invasion into forested areas (Silvério et al. 2013). An LAI of 3 represents approximately the inflection point of the logistic model after which small changes in LAI can quickly drive the system to one state or the other (Figure S1; Silvério et al. 2013). Because forests continue to lose tree cover after a fire (Brando et al. 2019), it is very likely that a transition to a low tree cover state occurs if canopy cover drops below the LAI value of 3 already immediately after a fire. Therefore, we can assume a high probability of grass invasion below this threshold. Indeed, other studies in regions of savanna-forest transitions have shown that an LAI of 3 marks the savanna-forest transition (Hoffmann et al. 2012, Cardoso et al. 2018). Because Silverio et al. (2013) found that this value corresponds to a probability of grass invasion of 30 % (Fig. S1), we took 30 % as the separation between low and high risk of grass invasion.

**Simulations**

The simulations were performed with fire-induced tree mortality rates predicted by CARLUC-Fire for 2010. The year 2010 was used to delineate the spatial distribution of our first loss term (drought effects, that essentially transfer part of the simulated live carbon stocks to litter material, increasing fuel loads leads to increasing fire intensity). We subsequently subtracted the loss term of CARLUC-Fire from the MODIS LAI values and obtained the probability of grass invasion using Equation S2. Thus, the simulation estimates the probability that a single fire event in a year of drought would promote grass invasion across the Amazon under current climate.

For the business-as-usual scenario we used climatic conditions from the CMIP5 models to adapt the biomass loss terms in CARLUC-Fire. Here, instead of directly using current LAI values from MODIS as pre-fire LAI, we summed these values by a correction term calculated based on simulated changes in ecosystem productivity under climate change. The correction term was calculated by (1) simulating present and future leaf biomass using the IBIS model; (2) multiplying these values by SLA to obtain LAI; and (3) calculating the difference between the estimated LAI for future and present condition mediated by differences in plant productivity resulting from climate change.

**Equation S1:** Relationship between MCWD and changes in biomass (Phillips et al 2009)

$$\Delta\ AGB = 0.3778 - 0.052 * \Delta MCWD \qquad \text{(Eq. S1)}$$

where AGB represents predicted losses in ABG and MCWD the maximum climatological water deficit.

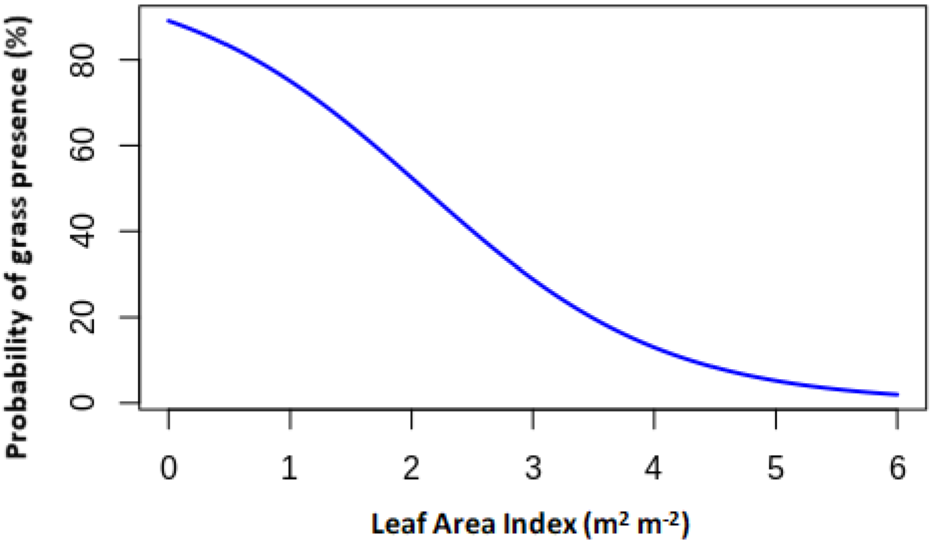


**Figure S1**- Probability of grass presence as a function of LAI from field measurements (Silvério et al. 2013; Eq. S2)

**Equation S2**: Probability of grass presence as a function of LAI

Prob. grass invasion = $100 * (1-exp(LAI+(-2.09))/(1 + exp(LAI+(-2.09))))$ (**Eq. S2**)

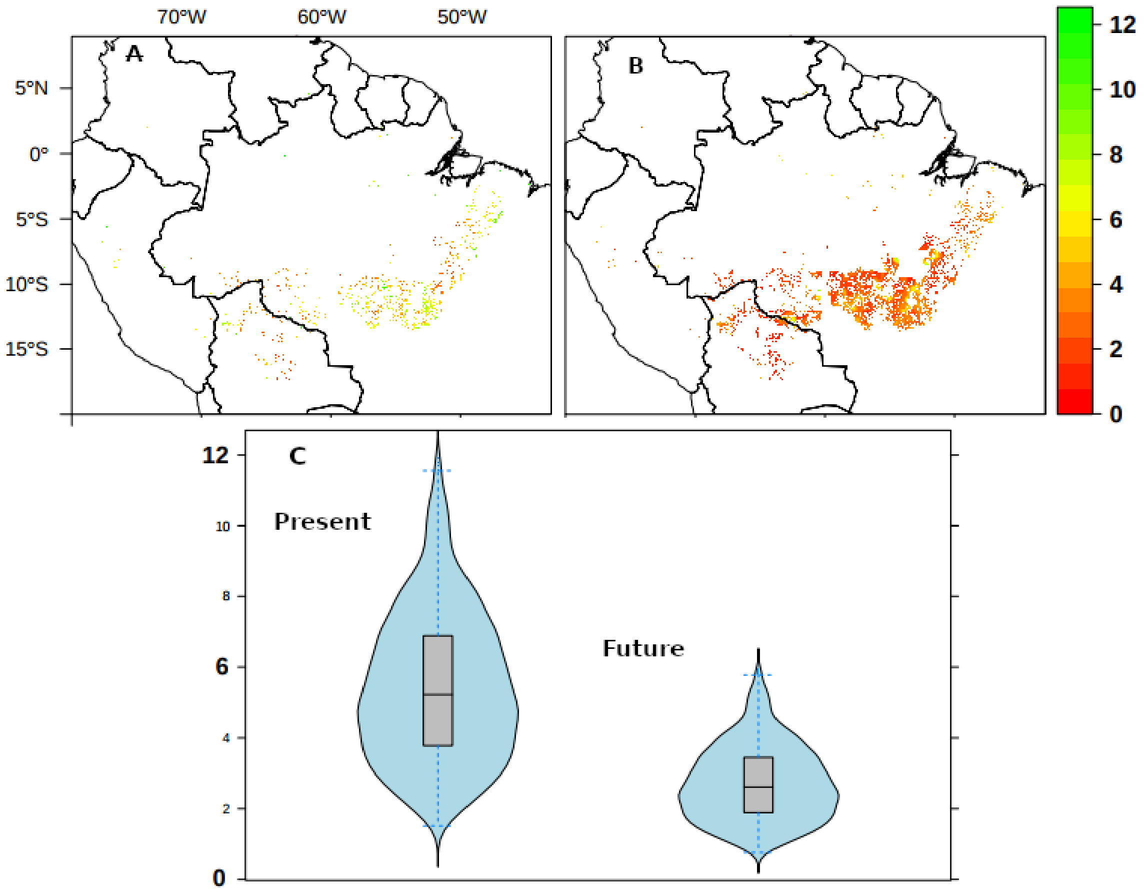


**Figure S2** - Fire return interval (FRI), defined as the number of years between two successive fire events, within regions with potential LAI < 3 after high-intensity fires. For present 2003-2016 (A) and future (B) with increased frequency of fire due grass-fire feedbacks. (C) The violin plots summarize FRI distributions, where the sides of each violin is a kernel density function

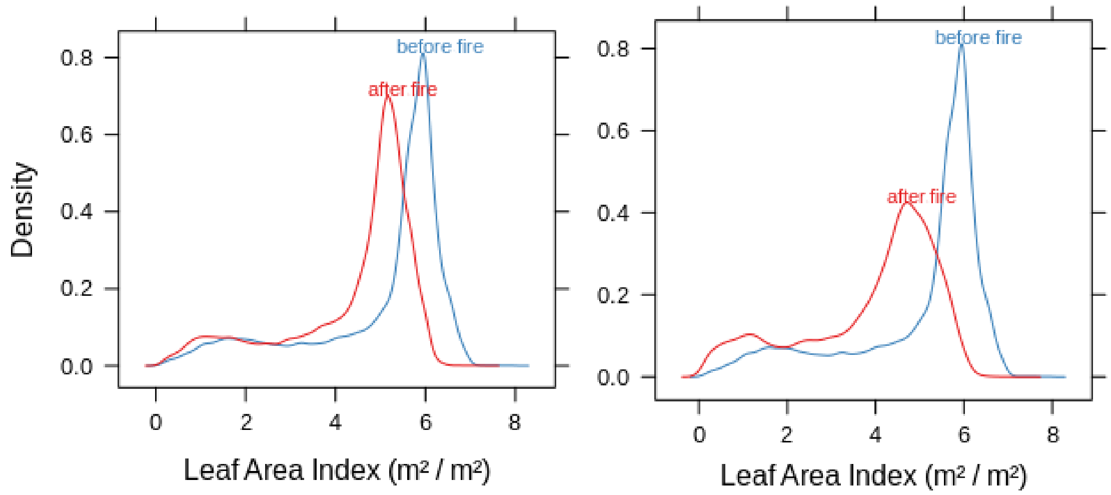


**Figure S3 -** Density distributions of Leaf Area Index (LAI) before and after a fire for the Amazon region under current (A) and future (B) climate scenarios.

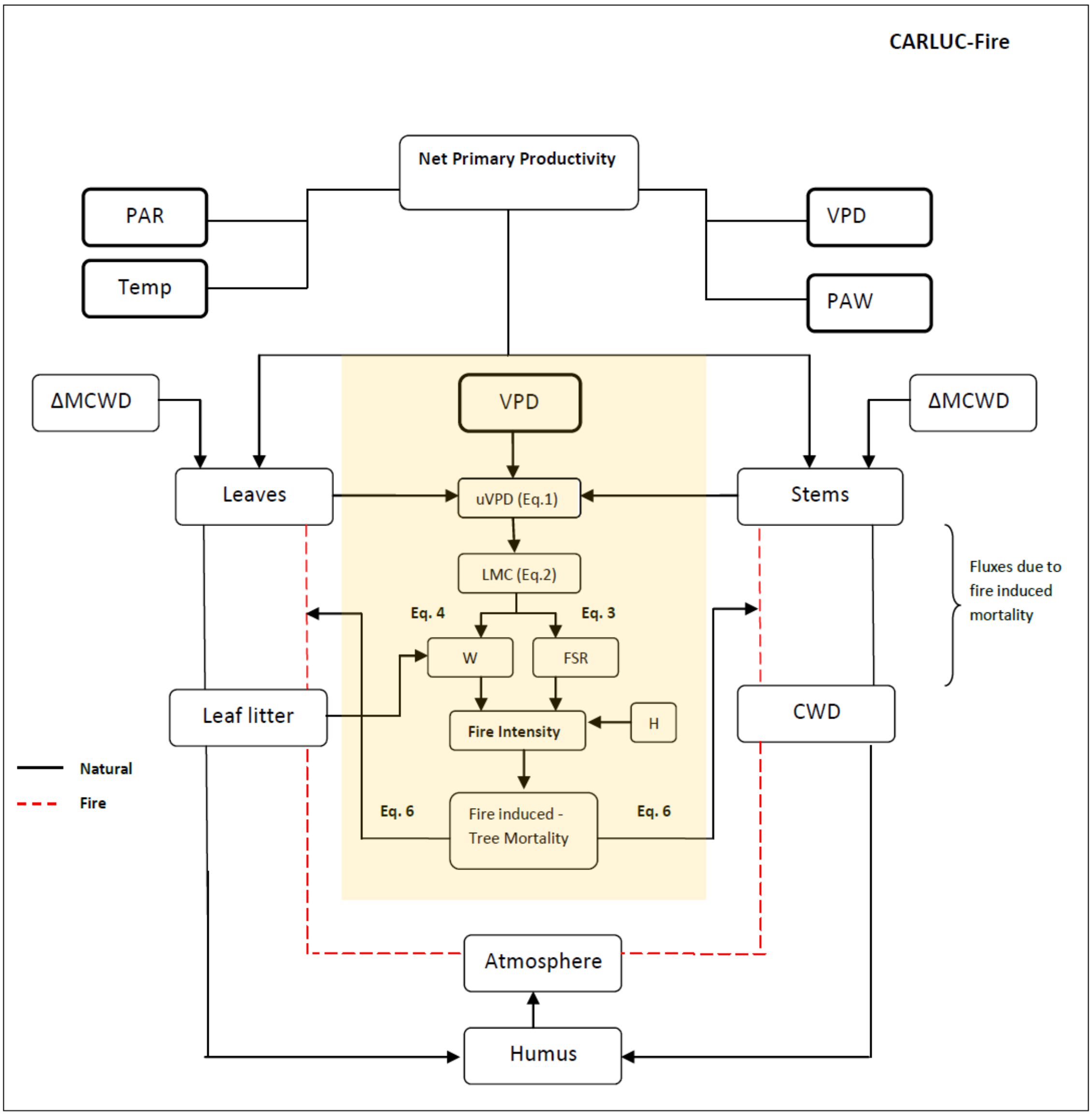


**Figure S4 -** Flowchart of the model focused in this study, in light orange the fire component

**Table S1:** Principal equations of CARLUC-Fire (De Faria et al. 2017)
Cstem: Carbon in stems. Cleaf: Carbon in leaves.
Cllstruc: Carbon in structural leaf litter. Unit for all carbon pools (kg/$m^2$)

| **Variable** | **Name** | **Unit** | **Equation** | **Eq. #** |
|---|---|---|---|---|
| uVPD | Inner Vapor Pressure Deficit | Kpa | 0.140494 - 0.006 * Cstem * 10 - 0.594074 * sqrt(Cleaf * 10 + 0.5) + 1.505 * sqrt( VPD + 0.5) | 1 |
| LMC | Litter Moisture Content | % | 80 * exp(-0.9 * uVPD) | 2 |
| FSR | Rate of Spread | m/min | 0.043 + 0.83 * exp(-0.107 * LMC) | 3 |
| W | Mass of fuel consumed by fire | Kg/$m^2$ | Fuel , LMC/me < 0.18<br>( 1.2 -0.62 * LMC/me) * Fuel, 0.18 ≤ LMC/me ≤ 0.73<br>(2.45 – 2.45 LMC/me) * Fuel , LMC/me > 0.73 | 4 |
| FI | Fire Intensity | kW/m | W * FSR * 18700[a] * 0.16 | 5 |
| Mort | Mortality[b] | Kg/$m^2$ | 1 / ( 1 + exp(2.45 - 0.002373 * FI ) | 6 |

[a] *The combustion heat (H), which is assumed to be constant at 18 700 kJ $kg^{-1}$ (Van Wagner 1973, Albini 1976); and mass of fuel consumed by fire (W), which is based on the assumption that the proportion of each dead fuel class that is consumed by fire decreases as a function of its moisture content relative to its moisture of extinction (me; following Peterson and Ryan 1986).*

[b] *fire-induced tree mortality (i.e. biomass turnover)*
